# Reduced visual and frontal cortex activation during visual working memory in grapheme-colour synaesthetes relative to young and older adults

**DOI:** 10.1101/444992

**Authors:** Gaby Pfeifer, Jamie Ward, Natasha Sigala

**Affiliations:** Department of Neuroscience, Brighton and Sussex Medical School, University of Sussex, Brighton BN1 9RR, UK; School of Psychology, University of Sussex, Brighton BN1 9RR, UK; Sackler Centre for Consciousness Science, University of Sussex, Brighton BN1 9RR, UK; Leeds School of Social Sciences, Leeds Beckett University, Leeds LS6 3QW, UK

**Keywords:** working memory, synaesthesia, healthy ageing, fMRI

## Abstract

The sensory recruitment model envisages visual working memory (VWM) as an emergent property that is encoded and maintained in sensory (visual) regions. The model implies that enhanced sensory-perceptual functions, as in synaesthesia, entail a dedicated VWM-system, showing reduced visual cortex activity as a result of neural specificity. By contrast, sensory-perceptual decline, as in old age, is expected to show enhanced visual cortex activity as a result of neural broadening. To test this model, young grapheme-colour synaesthetes, older adults and young controls engaged in a delayed pair-associative retrieval and a delayed matching-to-sample task, consisting of achromatic fractal stimuli that do not induce synaesthesia. While a previous analysis of this dataset (Pfeifer et al., 2016) has focused on cued retrieval and recognition of pair-associates (i.e. long-term memory), the current study focuses on visual working memory and considers, for the first time, the crucial delay period in which no visual stimuli are present, but working memory processes are engaged. Participants were trained to criterion and demonstrated comparable behavioural performance on VWM tasks. Whole-brain and region-of-interest-analyses revealed significantly lower activity in synaesthetes’ middle frontal gyrus and visual regions (cuneus, inferior temporal cortex) respectively, suggesting greater neural efficiency relative to young and older adults in both tasks. The results support the sensory recruitment model and can explain age and individual WM-differences based on neural specificity in visual cortex.

## Introduction

Visual working memory (VWM) refers to the transient mental rehearsal of visual stimuli that have been perceptually cued or retrieved from long-term memory, but are no longer present in the environment. VWM is supported by a distributed system, involving lateral regions of the prefrontal cortex (PFC), as well as parietal and occipital-temporal areas (Curtis and D’Esposito, 2003; D’Esposito, 2007; Postle, 2006; Ranganath, 2006; D’Esposito and Postle, 2015). However, the precise role of these brain regions has only been researched more recently. One WM model, dubbed the ‘sensory recruitment model’ (Serences et al., 2009; Lee and Baker 2016), envisages VWM as an emergent property from sensory regions as early as V1, which specifically code for feature and stimulus-specific information. An important characteristic of the model is the sustained representation of visual perceptual information along inferior occipito-temporal cortex, even after the perceptual stimulus has faded. Thus, the model suggests that VWM is maintained in the same posterior visual brain regions that are responsible for perceptual encoding. Early influential research has attributed WM-related stimulus representations to PFC rather than visual regions, e.g. (Baddeley and Hitch, 1974; Goldman-Rakic, 1990). A number of non-human primate studies reported spiking activity of single units during the delay-period of WM-tasks, which was taken as evidence for retained stimulus-specific information in PFC (Fuster 1973, Funahashi, Bruce et al. 1989, Goldman-Rakic 1995). However, converging findings with human participants and functional magnetic resonance imaging (fMRI) have since offered further insights into the specific frontal and posterior contributions to WM. For example, the distributed model of working memory envisages the PFC as an area exerting top-down control over posterior sensory regions. It converges with the sensory recruitment model on the notion that posterior sensory regions carry specific representational content (Postle, 2006; D’Esposito and Postle 2015; Lee and Baker 2016). Key support for the distributed and sensory recruitment model comes from studies using multi voxel pattern analysis (MVPA) that could discern the representational content in relevant frontal and occipito-temporal regions. Two studies (Christophel et al., 2012; Riggall and Postle, 2012) showed that although there was a sustained BOLD-response in frontal regions throughout the delay-period of a VWM task, decoding accuracy of the stimulus content was at chance-level. By contrast, no sustained BOLD-response could be detected within lateral occipito-temporal (Riggall and Postle, 2012) and early visual regions (Christophel et al., 2012), but decoding performance of the sub-threshold activity in these regions was significantly above chance-level. Other MVPA studies have recently reported high decoding accuracy of stimulus content in visual and frontal regions (Ester, Sprague et al. 2015; Sreenivasan, Vytlacil et al. 2014). However, only visual regions exhibited sensory-specific information while PFC exhibited non-sensory representations referring to higher-order task or goal orientations (Sreenivasan, Vytlacil et al. 2014). Together, these and other studies (Albers et al., 2013; Han et al., 2013; Ranganath et al., 2004) suggest that content-specific information of VWM is represented in occipito-temporal cortex, while the PFC appears to respond adaptively to task specific input (Sigala et al. 2008, Stokes, Kusunoki et al. 2013).

Concerning visual cortex activity, two studies have shown that the application of transcranial magnetic stimulation (TMS) over early visual cortex (V1 and V2) facilitated performance accuracy (Soto et al., 2012) and reduced response times (Cattaneo et al., 2009) during VWM tasks. These findings suggest that increased cortical excitability of visual regions, as induced via TMS-stimulation, can boost VWM. Here, we further tested this hypothesis by examining young grapheme-colour synaesthetes who show enhanced cortical excitability (Terhune et al., 2011) as well as enhanced sensitivity in early visual regions (Barnett et al., 2008), concomitant with superior performance on a range of cognitive tasks including WM (Rothen et al., 2012; Terhune et al., 2013). Grapheme-colour synaesthesia (in the following referred to as synaesthesia) is a stable perceptual phenomenon, found in about 1% of the population (Simner et al., 2006), whereby black letters, words, or digits (graphemes) are experienced as inherently coloured (e.g. the letter S may be perceived as green). Synaesthesia has a neurological basis, showing increased white matter connectivity in inferior temporal gyrus and superior parietal lobe (Rouw and Scholte, 2007), as well as increased grey-matter volume along the calcarine, lingual-and inferior temporal gyrus relative to controls (Banissy et al., 2012; Jancke et al., 2009; Rouw et al., 2011; Weiss and Fink, 2009). These anatomical differences are paralleled by functional differences in posterior brain regions and provide evidence of enhanced neural sensitivity in synaesthetes. Several studies were able to show activation in colour area V4 while synaesthetes processed black letters (Brang et al., 2010; Hubbard et al., 2005; van Leeuwen et al., 2011; Gould Van Praag et al., 2016); but see (Paulesu et al., 1995; Weiss et al., 2005; Rouw and Scholte, 2010; Hupe et al., 2011). Behaviourally, synaesthetes show a performance advantage over controls in WM for colour (Terhune et al., 2013) or in colour memory (Pritchard et al., 2013; Rothen and Meier, 2010; Rothen et al., 2012; Yaro and Ward, 2007), suggesting enhanced neural sensitivity in colour areas *per se*. Indeed, the synaesthetes’ frequent sensory experiences with colours following the secondary responses to words may sensitise colour areas in the brain and lead to enhanced colour processing (Banissy et al., 2009). However, the synaesthetes’ enhanced neural sensitivity goes beyond colour processing and is even found for stimuli that neither evoke a synaesthetic response, nor contain a perceptual colour. Perceptual processing of black pseudo-letters (that evoked no colour responses) yielded activity in the synaesthetes’ left inferior parietal lobe (IPL), which was not seen in controls (Sinke et al., 2012). Likewise, perceptual processing of abstract patterns with high spatial frequency and varying luminance contrast yielded enhanced early visually evoked potentials that were attributed to processing differences in primary visual cortex (Barnett et al., 2008). Although behavioural evidence for the enhanced processing account for non-synaesthesia inducing stimuli is mixed, a number of studies have shown an advantage of synaesthetes relative to controls in drawing abstract stimuli from memory (Gross et al., 2011; Rothen and Meier, 2010); but see (Yaro and Ward, 2007), and in recognising achromatic fractal images (Pfeifer et al., 2014; Ward et al., 2013).

In contrast to the enhanced visual sensitivity observed in synaesthetes, older individuals typically experience a loss of visual sensitivity. This has been explained as age-related neural broadening in ventral visual cortex (Park et al., 2004; 2012). Neural broadening is characterised by poorly differentiated neural responses to category selective stimuli (e.g. faces, houses, words). Category selective face, house and word areas in ventral visual cortex lose their sensitivity with age and respond broadly and less selectively across many stimuli (Park et al., 2004). Interestingly, the effects of age-related neural broadening are not limited to perceptual encoding, but have also been observed during visual imagery of category selective stimuli (Kalkstein et al., 2011). In fMRI, neural broadening is characterised by enlarged activation patterns, resulting from activation of many non-selective units across larger patches of cortex in response to different stimulus categories (Kok et al., 2012). By contrast, enhanced visual sensitivity and feature-selective responses are characterised by sparse and efficient encoding, resulting in smaller activation maps in fMRI. Evidence for age-related neural broadening was demonstrated by our previous fMRI analyses of the present dataset, which focused on associative memory (Pfeifer et al., 2016): Our group of older adults showed enhanced activation in visual cortex relative to synaesthetes and young adults during cued retrieval. The age-specific effect suggested that the increased visual cortex activation might have been the result of neural broadening and loss of visual sensitivity in older adults.

In the present study, we conducted a different analysis of the fMRI dataset from our previous study with young synaesthetes, and young and older non-synaesthetes (Pfeifer et al., 2016), in order to address a novel research question. The previous analysis focused on visual cues (the first item in a pair) and visual recognition (to determine if the stimulus was the corresponding item to the first). These involve memory processes involved, respectively, in generating internal representations in response to the cue and deciding whether that internal representation matches the presented one. Both, out of necessity, conflate visual perception and memory. In the current analysis, by focussing on the delay period (between cue and recognition), it is possible to study a different memory process (working memory maintenance of the internally generated visual representation) in the absence of visual perception. Specifically, we focused on the delay period of the delayed pair-associative (DPA) retrieval task and a delayed matching-to-sample (DMS) task. Both tasks involve stimulus 1 (cue), followed by delay, followed by stimulus 2 (recognition). While the delay period of the DPA-task required the maintenance of retrieved pair-associates from memory (high WM-load), the delay period of the DMS-task constituted a pure WM condition, simply requiring participants to hold a cued image in mind (low WM-load). The stimuli consisted of achromatic abstract fractal images, allowing us to test the enhanced processing hypothesis in synaesthetes for non-synaesthesia inducing stimuli, e.g. (Barnett et al., 2008; Rothen et al., 2012; Terhune et al., 2011; Yaro and Ward, 2007) and its relationship to VWM.

Insofar as synaesthetes show enhanced neural sensitivity in feature-selective and non-selective regions in occipito-temporal cortex (Pfeifer et al., 2016; Barnett et al., 2008; Brang et al., 2010; Hubbard et al., 2005), we predicted activation differences in these regions relative to young and older adults during VWM maintenance. Specifically, older adults might show greater activity than synaesthetes in inferior temporal regions as a result of age-related neural broadening (Park et al., 2012). Neural broadening opposes the neural specificity found in synaesthetes in that feature-selective neurons lose their selectivity (e.g. the fusiform face area in response to faces) and code for a variety of other visual stimuli. Consequently, age-related neural broadening in inferior temporal cortex would yield increased BOLD-responses in fMRI relative to synaesthetes, and possibly young adults.

We further expected group differences in early visual regions. A key finding in our previous report (Pfeifer et al., 2016) was that the present group of participants showed activation differences in early visual cortex during visual associative memory. Specifically, we found enhanced activation in synaesthetes’ early visual cortex relative to young and older adults during the recognition stage of the DPA and DMS tasks, reflecting enhanced sensitivity to external, behaviourally relevant stimuli (cf. Terhune et al., 2011; Barnett et al., 2008). By contrast, activation was reduced in synaesthetes’ early visual cortex relative to young and older adults during cued retrieval, reflecting selective coding of internally represented associative memories (cf. Kalkstein et al., 2011). The present analyses focus on VWM, which requires internal, mental representations of visual stimuli. Hence, we predicted lower activation in synaesthetes’ visual cortex relative to young and older adults, based on our previous findings. Alternatively, our whole-brain analyses might not detect a group difference in occipito-temporal regions, given that the content-specificity of maintained stimuli in posterior visual areas is often not accompanied by a sustained BOLD-response (Christophel et al., 2012; Riggall and Postle, 2012).

Prefrontal cortex activity was expected to be enhanced in older adults as a compensatory strategy for neuronal dedifferentiation in occipital-temporal cortex, described as the posterior-to-anterior shift (Davis, Dennis et al. 2008). Finally, the group differences were expected to be modulated by task difficulty. We predicted activation differences between the two WM-tasks, based on findings of differential neural activity for different types of information maintained in WM (Curtis et al., 2004; D’Esposito, 2007; Ranganath et al., 2004).

## Materials and Methods

### Participants

Nineteen young adults (8 female; age range = 21 – 32 years; M = 24.32), nineteen older adults (11 female; age range = 59 – 81 years; M = 66.21), and nineteen young grapheme-colour synaesthetes (15 female; age range = 19 – 33 years; M = 23.00) took part in the experiment, which was reviewed and approved by the Brighton and Sussex Medical School Research Governance and Ethics Committee. The same participants took part in our previously reported fMRI study that focused on visual associative memory (Pfeifer et al., 2016). The participants had no history of psychiatric or neurological diseases. The average number of years of formal education for young adults was M = 16.95 (SD = 1.68), for older adults M = 13.95 (SD = 3.32), and for the synaesthetes M = 16.74 (SD = 2.11). The groups differed in the number of years of education (*F*[2,54] = 8.717; *p* = 0.001). Tukey post hoc comparisons showed that the difference was significant between young and older adults (*p* = 0.001), between synaesthetes and older adults (*p* = 0.003), but not between young adults and synaesthetes (*p* = 0.963). Screening for cognitive impairment was carried out for all but 5 young adults, using the Mini Mental State Examination [MMSE; (Folstein et al., 1975)]. All participants performed comparably on the MMSE, *F*[2,51] = 2.11; *p* = 0.131, with high average scores across the 14 young adults (M = 28.93; SD = 0.93), 19 older adults (M = 28.15; SD = 1.46), and 19 synaesthetes (M = 28.89; SD = 1.37). Synaesthetes were recruited from the University of Sussex and via the UK Synaesthesia association website www.uksynaesthesia.com. All synaesthetes reported seeing colours in response to letters or digits. To verify Synaesthesia, we used the ‘Synesthesia battery’ (Eagleman et al., 2007), available on www.synesthete.org, and the adapted cut-off score of 1.43 (Rothen et al., 2013). Using this battery, a mean score of M = 0.81 (SD = 0.28; range = 0.38 – 1.39) was obtained across our group of synaesthetes, which is consistent with synaesthesia.

### Experimental design and Stimuli

The fMRI protocol consisted of a delayed pair-associative (DPA) retrieval task and a delayed matching-to-sample (DMS) task. The DPA-task was always presented first in order to avoid retroactive interference effects on participants’ associative memory.

#### DPA-task

For the DPA-task, we selected eight pair-associates (black-and-white fractal images) from a pool of 16 pairs. The eight pair-associates were divided into four visually similar and four visually dissimilar pairs to create a low and high memory load condition, respectively. The visual similarity of all pair-associates was rated by an independent group of 20 participants. This has been described in more detail in Pfeifer et al. 2014. Participants gave their ratings on a 5-point Likert scale (Likert, 1932), where a rating of 1 indicated no visual similarity and a rating of 5 indicated high visual similarity between pairs. Based on the mean-ratings, we selected the 4 most dissimilar and the 4 most similar pairs respectively, representing high and low memory load conditions (example pairs illustrated in Figure 1). A Wilcoxon signed-rank test demonstrated that the 4 selected similar pairs were rated significantly higher in visual similarity (M = 3.87; SD = 0.38) compared to the four selected dissimilar pairs (M = 1.31; SD = 0.20); significance Z = -2.521; p = 0.012 (two-tailed). We used an event-related design, during which each of the selected pairs was randomly presented eight times, amounting to a total of 32 similar and 32 dissimilar pairs. The cue and target images were presented interchangeably throughout the task. On 62.5 % of the trials, the cue pictures were followed by a matching target, constituting 40 match-trials and 24 non-match trials. In this sense, lure stimuli were non-matching images from the same set of the 8 pair-associates rather than trial unique stimuli. Using recombinations of same-set stimuli constitutes a more powerful test of associative memory, requiring participants to retrieve the intact combination of pair-associates out of equally familiar stimuli rather than rejecting lures on the basis of their novelty (Mayes et al., 2007).

#### DMS-task

For the DMS-task, we chose an independent set of 8 individual black-and-white fractal images. The DMS-task constituted our third condition to be compared against the low and high memory load condition of the DPA-task. We used an event-related design, consisting of a pseudo-random presentation of 32 individual fractal images, with each of the selected 8 images shown 4 times. On 62.5 % of the trials the cue pictures were followed by a matching target, constituting 20 match-trials and 12 non-match trials. Lure stimuli were non-matching images from the same set of the 8 fractals rather than trial unique stimuli. Across the DPA and DMS-task, the minimum trial distance between match and non-match trials was one (i.e. a match trial could immediately follow a non-match trial and vice versa), and the maximum trial distance was five (i.e. a non-match trial could follow four presentations of match-trials).

The behavioural data of the DPA and DMS-task were initially analysed using a 3 × 3 (group × condition: low and high memory load, DMS) mixed ANOVA. Since we did not find a behavioural effect between the DMS-task and the low memory load condition of the DPA-task, we only compared the high memory load condition of the DPA-task against DMS in our fMRI analysis (details provided under fMRI analyses and Table 1).

**Figure 1.**
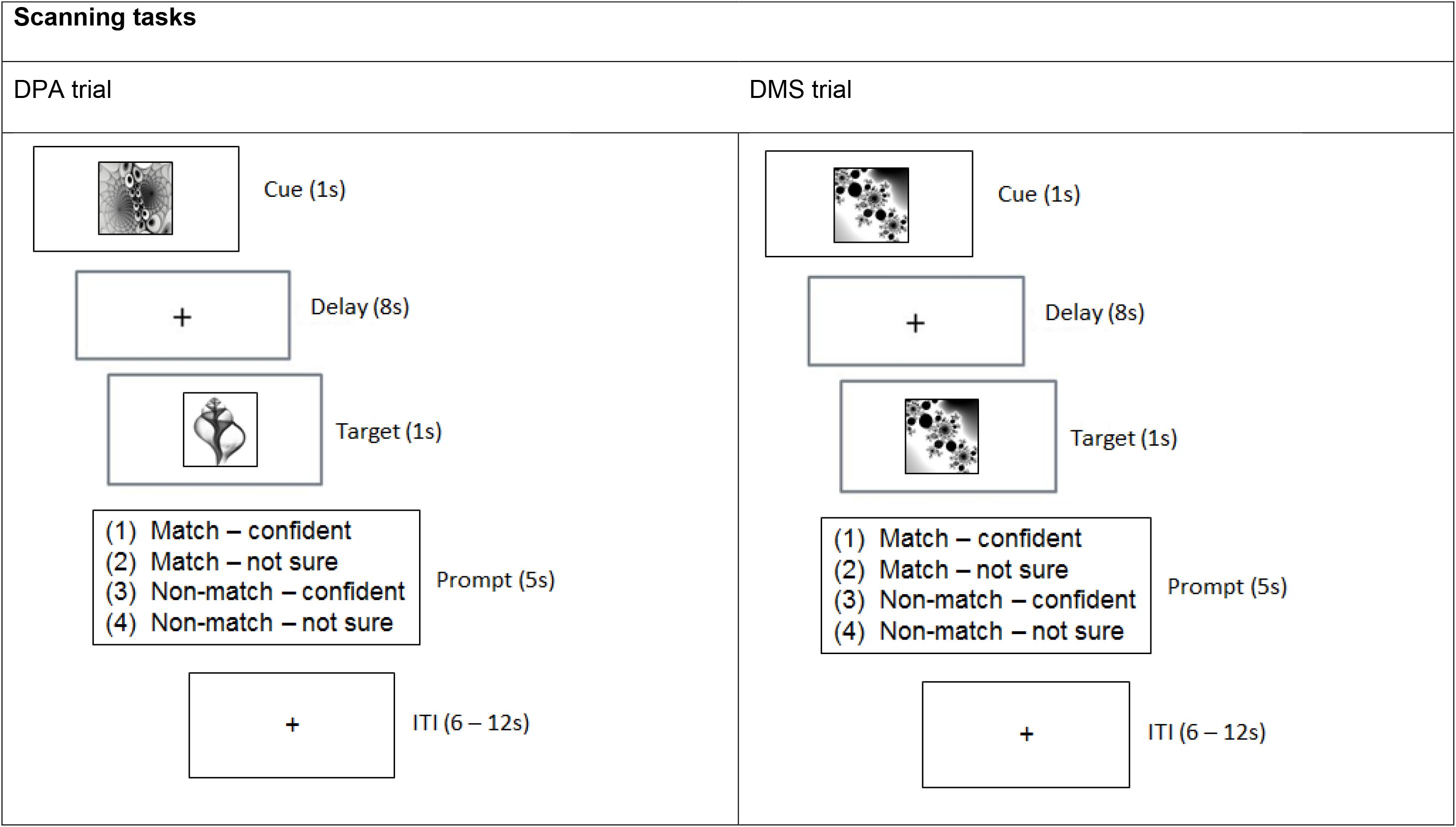
Experimental design. The scanning tasks involved two trial types, DPA and DMS. DPA trials required participants to retrieve a cue’s matching pair-associate and hold it in mind over an 8 second delay. DMS trials required participants to hold the cue in mind over an 8 second delay. Upon target presentation, participants were asked to decide whether the target was a match or non-match (in DPA and DMS trials) and give their responses within a 5 second time window (Prompt). ITI = Interstimulus interval; s = second.

### Procedure

Prior to scanning, participants were trained on the fractal pair-associates of the DPA-task. The task began with the sequential presentation of eight pair-associates at the centre of a computer screen for 4s, and participants were instructed to remember the correct association of the pairs for a subsequent memory test. The presentation was followed by a four alternative forced-choice task, in which participants had to choose one of four possible target pictures from the bottom of the screen to match the cue picture at the top of the screen. The pictures stayed on screen until a response was recorded. Each response was followed by a 3s presentation of visual feedback, indicating whether or not the matching target had been identified correctly (green tick or red cross respectively). Participants performed the task until they reached an 87.5% learning criterion. A minimum of 2 Runs was required in the learning phase. Cue and target shapes of all pair-associates were presented interchangeably during learning: a stimulus that had been presented as the cue in one Run became the target in the following Run. Stimuli were delivered using Presentation^®^ 14.9 (Neurobiobehavioral Systems, Inc., Berkeley, CA).

#### DPA and DMS

Following the associative learning task, participants were familiarised with the DPA and DMS-task prior to scanning. During scanning, an identical trial structure was used across the DPA and DMS-task (Figure 1). During the cue-period (1s) of the DPA-task, participants were asked to use the cue to retrieve the matching target (*associative retrieval*). During the cue-period (1s) of the DMS-task, participants were asked to build up a mental image of the cue. The delay period (8s) required participants to either hold the retrieved picture in mind (DPA-task), or to hold the cue image in mind (DMS-task). Finally, the target presentation (1s) in the DPA-task comprised the *associative recognition* stage, where participants were asked to recognise the target as the matching or non-matching pair-associate. The target presentation (1s) of the DMS-task required participants to judge whether the target was the identical image to the cue. Following target presentation in both tasks, a response window appeared for 5 seconds, during which participants were asked to press 1 of 4 buttons, providing combined decisions about the target (match/non-match) and self-rated confidence (confident/not sure). The button-presses were followed by variable intertrial intervals (ITI) of 6 – 12 s.

### Data acquisition

Imaging data were collected using a 1.5 Tesla MRI scanner (Siemens Magnetom Avanto) with a 32-channel phased-array head coil, tuned to 66.6 MHz. Visual stimuli were presented on an in-bore rear projection screen, at a viewing distance of approximately 45 cm, subtending 5 degrees of visual angle. Stimuli were delivered using Cogent2000 v1.32 running under MATLAB R2006b (The MathWorks, Inc., Natick, MA). Time-course series of the two runs were acquired using a T2*-weighted echo planar imaging (EPI) sequence, obtaining 644 volumes during the DPA-task, and 324 volumes during the DMS-task. Each volume consisted of 35 axial slices oriented in parallel to the AC-PC line, and covering the whole brain. Slices were acquired bottom-up in the interleaved mode. The following functional imaging parameters were used: TR=2620ms, TE=42ms, flip angle 90°, matrix= 64×64, FoV=192×192mm, slice thickness=3.0mm with a 20 % gap, resulting in 3.0mm isotropic voxels. To aid distortion correction, corresponding phase and magnitude field maps were acquired with a TR=513ms, TE1=5.78ms, TE2=10.54ms, flip angle 60°. A whole-brain, high-resolution T1-weighted 3D structural image was obtained using a magnetisation-prepared gradient-echo sequence, consisting of 192 contiguous axial slices (TR=1160ms, TE=4.24ms, flip angle 15°, matrix = 256×256, FoV= 230×230mm, 0.9mm isotropic voxel size). The T1-weighted image was used as an anatomical reference for each participant’s functional data.

### fMRI analyses

We used SPM8 (Wellcome Trust Centre for Neuroimaging, UCL, London, UK; www.fil.ion.ucl.ac.uk\spm) running under MATLAB R2013a for data preprocessing and statistical analyses. Preprocessing of functional images was carried out for each task separately, including slice-time correction to the middle slice, spatial realignment to the first image, and unwarping using the acquired field maps. The T1-weighted structural image was co-registered to the mean functional image and subsequently segmented to obtain normalisation parameters based on the standard MNI template. The segmentation parameters were used to transform each subject’s functional images and the bias-corrected structural image into MNI space. Voxel sizes of the functional and structural images were retained during normalisation, and the normalised functional images were spatially smoothed using an 8mm Gaussian kernel (full-width-half-maximum). Statistical analyses were performed using the General Linear Model. For the single subject analysis, the DPA and DMS-task were entered as separate sessions into the model. Across tasks, we specified regressors associated with the cue, delay, target and baseline (ITI) period. All regressors of interest contained only accurate and confident responses. The specific trial count for all accurate and confident responses included in the single subject analyses is detailed in Table 1. Modelling of regressors was similar across the DPA and DMS-task, given the identical trial structure: For each regressor representing a cue and target-period, activation was modelled using a boxcar function, starting at onset and lasting for 1 second. For the DPA task, two regressors were modelled for the cue, delay and target periods representing the retrieval of similar and dissimilar pair-associates, respectively, while there was only one condition/regressor representing the cue, delay and target periods for the DMS task [results relating to the cue and target periods were reported in Pfeifer et al. (2016)]. The delay-period was the main regressor of interest for the present study. The delay was modelled to start 3 seconds after delay-onset for a duration of 5 seconds, until the end of the delay-period. This was done to avoid capturing any residual activity pertaining to the cue-period, but instead explaining a largely unique source of variance pertaining to delay-period activity (Rissman et al., 2004). Baseline regressors were modelled to start 3 seconds after prompt-offset and lasted for 5 seconds. In instances of short ITIs of 6 seconds, baseline regressors were modelled to start 3 seconds after prompt-offset and lasted for 3 seconds. The baseline duration was chosen to match the duration of the delay-period to serve as a contrast for delay-period activity. Regressors of no interest included the prompt (containing participant’s button presses), a nuisance regressor (containing all misses, false alarms, non-confident responses, empty key responses) and six motion regressors. All regressors were convolved with a canonical hemodynamic response function available in SPM8 (Friston et al., 1998). A high-pass filter was applied with a period of 128 seconds to remove low-frequency signals relating to scanner drift and/or physiological noise. Two t-contrasts were computed comparing the two types of WM against Baseline: DPA Delay > DPA Baseline (DPAd > DPAb) and DMS Delay > DMS Baseline (DMSd > DMSb). DPA-related contrast images only included trials of the high memory load condition (i.e. dissimilar pair-associates) for the strongest comparison of WM for retrieved pair-associates versus WM for cued singletons.

**Table 1.** Trial count of all accurate responses of the DMS and DPA-task, separated by confidence ratings.

| **DMS-task** | **Hits** |  | **Hits Total** | **Correct Rejections** |  | **Correct Rejections Total** | **Grand Total** |
| --- | --- | --- | --- | --- | --- | --- | --- |
|  | **Confident** | **Not sure** |  | **Confident** | **Not sure** |  |  |
| **Young Adults** | 395 | 4 | 399 | 237 | 6 | 243 | 642 |
| **Older Adults** | 383 | 4 | 387 | 234 | 0 | 234 | 621 |
| **Synaesthetes** | 387 | 5 | 392 | 229 | 5 | 234 | 626 |

| DPA-task | Dissimilar |  |  |  | Dissimilar Total | Similar |  |  |  | Similar Total | Grand Total |
| --- | --- | --- | --- | --- | --- | --- | --- | --- | --- | --- | --- |
|  | Hits |  | Correct Rejections |  |  | Hits |  | Correct Rejections |  |  |  |
|  | Confident | Not sure | Confident | Not sure |  | Confident | Not sure | Confident | Not sure |  |  |
| Young Adults | 362 | 22 | 215 | 4 | 603 | 384 | 15 | 250 | 14 | 663 | 1266 |
| Older Adults | 257 | 32 | 189 | 1 | 479 | 373 | 2 | 252 | 2 | 629 | 1108 |
| Synaesthetes | 328 | 45 | 199 | 8 | 580 | 391 | 2 | 247 | 7 | 647 | 1227 |

Trial count for all accurate responses of the DMS-task (top) and DPA-task (bottom). Gray-shaded columns highlight trials that were included in the fMRI analyses. These trials included all confident, accurate responses (averaged across Hits and Correct rejections). For the DPA-task, only dissimilar accurate and confident trials were included in the fMRI analyses to achieve the strongest comparison of WM for retrieved pair-associates (DPA) versus WM for cued singletons (DMS).

#### Grey matter volume

Given that we compared a group of 19 older adults against 38 younger adults (19 synaesthetes and 19 controls) and had an unequal gender distribution across our 57 participants (male: N = 23; female: N = 34), we calculated each participants’ total grey matter (GM) volume in millilitre (ml). This value was subsequently entered as a covariate in all second-level fMRI analyses to implicitly account for age- (Lemaitre et al., 2005; Raz et al., 2005) and gender-related (Luders et al., 2002) GM volume differences. Total GM volume was calculated from the subject-specific GM masks in native space, which were obtained following the segmentation of participants’ high resolution structural T1 images.

#### Second-level analyses

To analyse brain activity associated with WM maintenance of retrieved pair-associates (DPA-task) and of cued singletons (DMS-task), the results of the single-subject analyses were taken to group-level. Using a 3 (group) × 2 (task) factorial ANOVA, we examined task, group, and group by task interaction effects using the contrast images DPAd>DPAb and DMSd>DMSb. We created exclusive masks for the average activity across DPA and DMS, as well as for the average activity of the DPA and DMS-task separately, using a t-contrast across groups and a lenient threshold of p < 0.01 (uncorrected). Task, group and interaction effects were computed using an F-contrast. They were inclusively masked with the respective average task activity and suprathresholded at *p* < 0.001 (uncorrected), k= 5 voxels. Thus, the masking ensured that *a)* group and interaction effects showed significant activations above zero within task-related regions and *b)* activity was reported at a more stringent threshold, as voxels had to survive the thresholds of the task effect as well as the group effect (Daselaar et al., 2010). To further explore brain areas showing group and interaction effects, we extracted the percent signal change of each mean cluster activity using the rfx-plot toolbox (Gläscher 2009). For brain areas showing a group difference, we estimated the trial-averaged BOLD signal change relative to cue-onset in second increments and plotted the time course for the average activity across DMS and DPA. For brain areas showing a significant group by task interaction, we presented the trial-averaged responses of all groups as the mean percent signal change relative to our modelled delay onset (starting 3 seconds into the delay).

## Results

### Behavioural results

Accuracy was high in both tasks and comparable across groups (see Table 2). A 3 × 3 mixed ANOVA with group and task as factors yielded no significant main effect of group, *F*[2,54] = 2.071, p = 0.136, η_p_^2^ = 0.071. A highly significant main effect of task (*F*[2,108] = 29.119, p < 0.001, η_p_^2^ = 0.350) suggested that the retrieval of dissimilar pairs was more demanding than the retrieval of similar pairs and the DMS-task (p < 0.001 for both pairwise comparisons, respectively). No difference was found between the retrieval of similar pairs and the DMS-task (p < 0.290). We also found a significant interaction between group and task, *F*[4,108] = 6.827, p < 0.001, η_p_^2^ = 0.202. Tests of within-subject contrasts showed that the difference was found between the similar and dissimilar retrieval condition (*F*[2,54] = 6.173, *p =* 0.004, *ηp^2^* = 0.186), and was driven by poorer performance of older versus young adults (parameter estimates: t = 3.214; p = 0.002).

**Table 2.** Mean and standard error of the percent Accuracy (Hits and Correct Rejections) in the DPA and DMS-task (N = 19 in each group).

| **Hit-rate (Task)** | **Young adults**<br>Mean (SE) | **Older adults**<br>Mean (SE) | **Synaesthetes**<br>Mean (SE) |
| --- | --- | --- | --- |
| Accuracy (DPA, similar pairs) | 93.46 (1.73) | 96.81 (0.71) | 96.81 (1.46) |
| Accuracy (DPA, dissimilar pairs) | 96.69 (2.16) | 73.85 (5.26) | 84.55 (4.39) |
| Accuracy (DMS) | 96.22 (1.21) | 96.38 (1.25) | 93.87 (1.37) |

### fMRI results

#### Main effect of Task: Brain activity is mediated by working memory demands

We discovered a main effect of task in the superior medial prefrontal cortex (PFC), inferior frontal and middle orbital gyrus, the insula and midline regions (including the anterior, middle, posterior cingulate gyrus and precuneus), the pre and post central gyrus, inferior parietal regions (supramarginal and angular gyrus), middle and superior temporal gyrus, inferior and middle occipital gyrus and the cerebellum. Post hoc tests revealed that DMS-related WM (t-contrast: DMSd>DMSb **>** DPAd>DPAb) activated the medial PFC, lateral temporal regions and inferior parietal cortex, as would be expected from a visual working memory task. By contrast, DPA-related WM (DPAd>DPAb **>** DMSd>DMSb) yielded greater activity in left lateral PFC and superior parietal cortex, consistent with associative retrieval (Table 3).

**Table 3.**
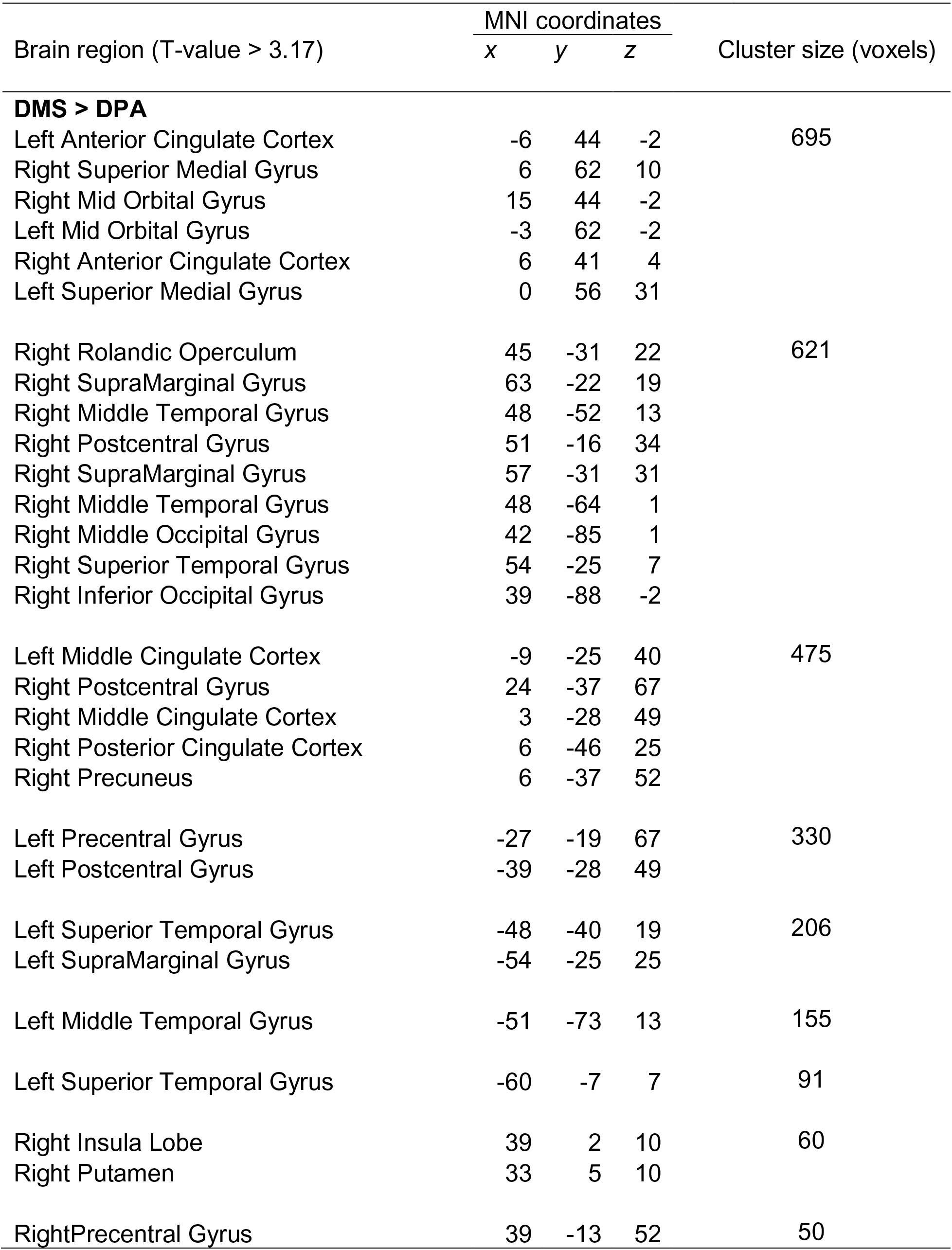
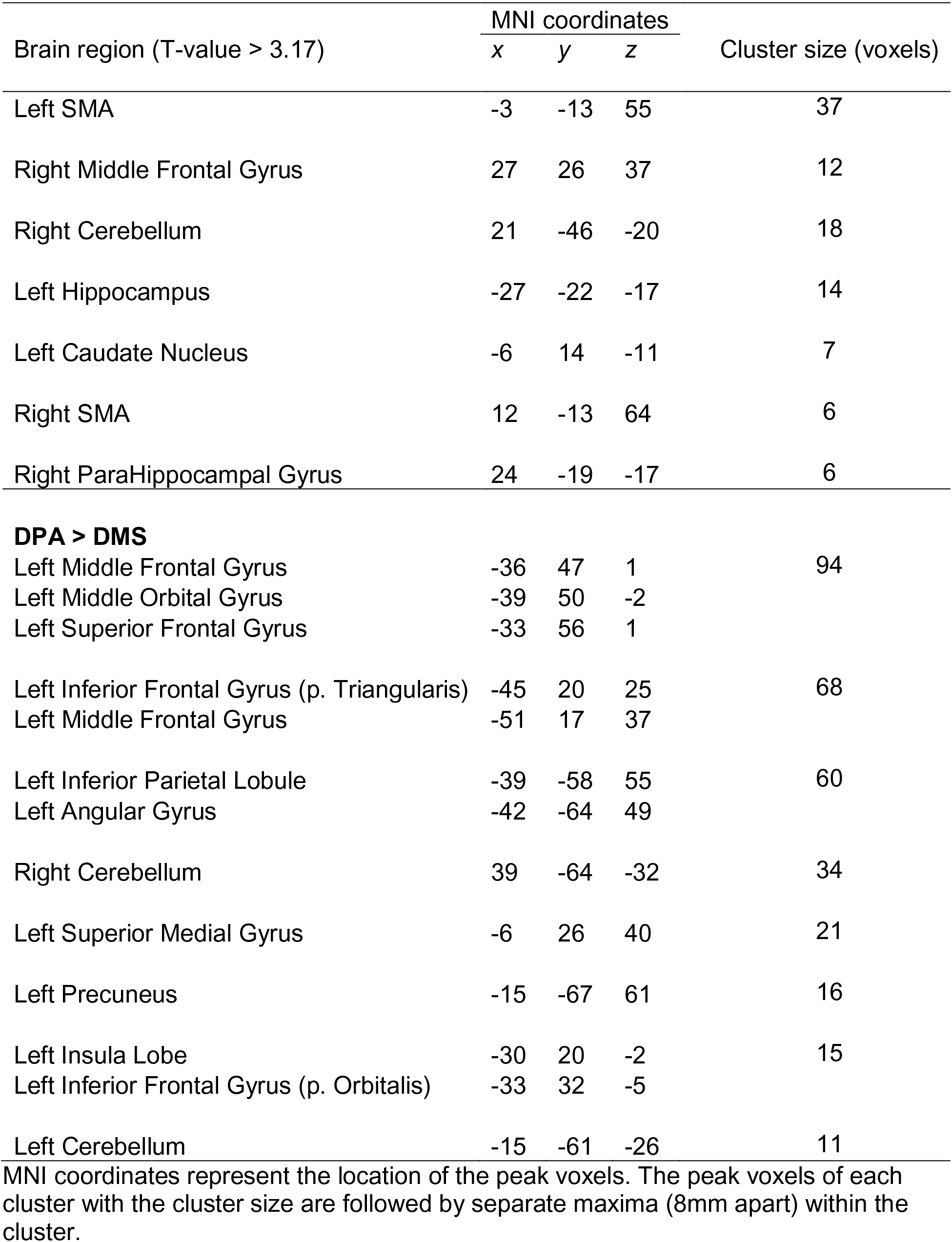
Regions showing activation differences between the DMS and DPA task, obtained from t-contrasts of the 3 × 2 (group × task) mixed ANOVA.

#### Main effect of group: Synaesthetes show reduced responses in visual, parietal and frontal regions during visual working memory

We found a significant main effect of group on WM maintenance in occipital, inferior temporal and frontal regions (Fig.2). Synaesthetes exhibited lower activations relative to the other two groups in all brain regions (Fig.2). The reduction in percent signal change in the synaesthetes’ occipital-temporal regions was specific to the delay period of the WM-tasks and was not seen during the cue and target stages of the task (Fig.2 A-D, G). Next, we computed t-contrasts to test for pair-wise group differences. We found a synaesthesia-specific effect in the inferior occipital and inferior temporal gyrus (averaged across DPA and DMS), with young adults showing higher activity than synaesthetes. No significant effect was found for the opposite contrast (synaesthetes > young), suggesting enhanced neural specificity in synaesthetes’ early visual regions. Older adults’ WM-maintenance was associated with higher activity relative to synaesthetes and young adults in occipital, parietal and frontal regions (Table 4). Notably, older adults showed more widespread activation differences relative to synaesthetes than relative to young adults, encompassing inferior temporal, fusiform and frontal regions. The opposite contrasts (young > old; synaesthetes > old) did not show an effect.

**Figure 2.**
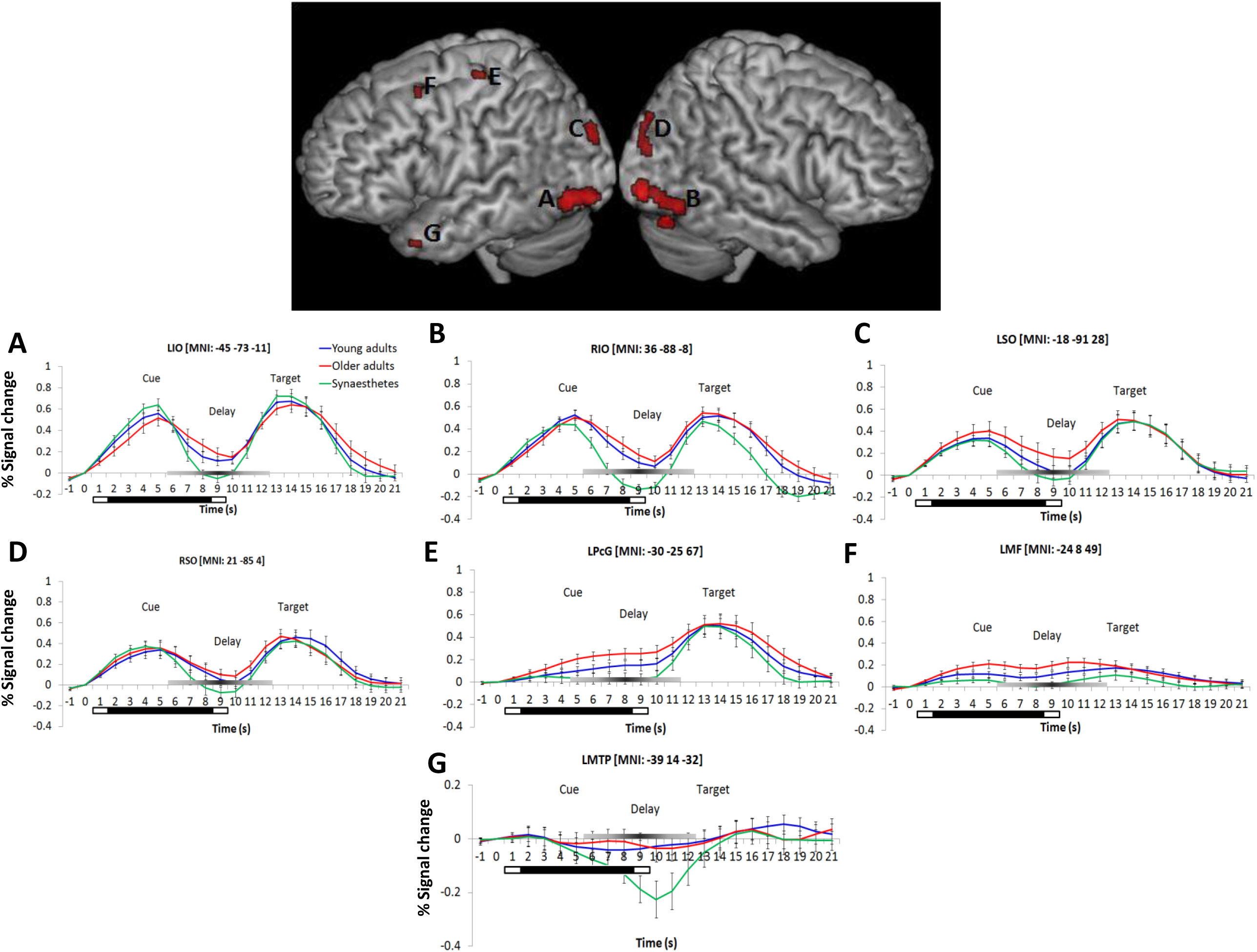
Left and right-hemispheric regions exhibiting a significant main effect of group for the delay period (averaged across DPA and DMS). Parametric maps are rendered on the individual subjects’ brain available in MRIcron. Regions included the bilateral inferior temporal gyrus (**A, B**), bilateral superior occipital gyrus (**C, D**), the left precentral gyrus (BA6; **E**), left middle frontal gyrus (BA9; **F**), and the left medial temporal pole (BA38; **G**). Further group differences were found in the cuneus, calcarine, lingual and fusiform gyrus, as well as in the left precuneus (not illustrated). Bottom: Peristimulus time histograms (PSTH). Trial-averaged responses for all groups were rescaled to the percent signal change of each mean cluster activity relative to cue-onset. Error bars represent the standard error of the mean. Reduced signal strength was found in synaesthetes relative to young and older adults during the delay period in regions **A-G**. The bars below each x-axis indicate the timing of cue and target stimulus presentation (white bar) and the delay period (black bar). A grey-scaled gradient bar above each x-axis depicts the expected peak of the BOLD response for WM-related activity, assuming a 4-6 second peak latency of the hemodynamic response. LIO = Left Inferior Occipital Gyrus, RIO = Right Inferior Occipital Gyrus, LSO = Left Superior Occipital Gyrus, RSO = Right Superior Occipital Gyrus, LPcG = Left Precentral Gyrus, LMF = Left Middle Frontal Gyrus, LMTP = Left Medial Temporal Pole. Regions of each PSTH are denoted with their cluster peak in MNI.

**Table 4.** Regions showing activation differences between young adults, older adults and synaesthetes, obtained from t-contrasts of the 3 × 2 (group × task) mixed ANOVA.

| Brain region (T-value > 3.17) | MNI coordinates |  |  | Cluster size (voxels) |
| --- | --- | --- | --- | --- |
|  | x | y | z |  |
| **Young > Old** |  |  |  |  |
| no effect |  |  |  |  |
| **Old > Young** |  |  |  |  |
| Left Cuneus | -12 | -88 | 28 | 24 |
| Left Superior Occipital Gyrus | -18 | -91 | 25 |  |
| Right Middle Cingulate Gyrus | 9 | -40 | 43 | 21 |
| Right Precuneus | 6 | -43 | 49 |  |
| Right Superior Occipital Gyrus | 18 | -85 | 34 | 15 |
| Right Cuneus | 12 | -82 | 34 |  |
| Left Precuneus | -12 | -43 | 49 | 11 |
| Left Middle Frontal Gyrus | -24 | 8 | 49 | 9 |
| Right Superior Occipital Gyrus | 21 | -88 | 25 | 7 |
| Left Precuneus | -3 | -58 | 43 | 6 |
| **Young > Synaesthetes** |  |  |  |  |
| Left Inferior Occipital Gyrus | -42 | -67 | -5 | 48 |
| Left Inferior Occipital Gyrus | -30 | -88 | -11 | 22 |
| Right Inferior Occipital Gyrus | 36 | -88 | -8 | 16 |
| Left Cerebellum | -33 | -73 | -20 | 8 |
| Right Inferior Temporal Gyrus | 54 | -64 | -11 | 7 |
| **Synaesthetes > Young** |  |  |  |  |
| no effect |  |  |  |  |

**Table 4 continued.**
| Brain region (T-value > 3.17) | MNI coordinates |  |  | Cluster size (voxels) |
| --- | --- | --- | --- | --- |
|  | x | y | z |  |
| **Synaesthetes > Old** |  |  |  |  |
| no effect |  |  |  |  |
| **Old > Synaesthetes** |  |  |  |  |
| Left Inferior Occipital Gyrus | -45 | -73 | -11 | 163 |
| Left Fusiform Gyrus | -30 | -79 | -17 |  |
| Left Lingual Gyrus | -27 | -88 | -14 |  |
| Left Superior Occipital Gyrus | -18 | -91 | 28 | 144 |
| Left Cuneus | -12 | -88 | 28 |  |
| Left Calcarine Gyrus | -12 | -85 | 13 |  |
| Right Superior Occipital Gyrus | 21 | -85 | 34 | 121 |
| Right Middle Occipital Gyrus | 39 | -85 | 22 |  |
| Right Cuneus | 9 | -79 | 34 |  |
| Right Inferior Occipital Gyrus | 36 | -88 | -8 | 75 |
| Right Inferior Temporal Gyrus | 51 | -64 | -11 |  |
| Right Cerebellum | 15 | -76 | -20 | 72 |
| Left Cerebellum | -6 | -76 | -14 |  |
| Right Calcarine Gyrus | 18 | -76 | 16 | 46 |
| Right Cuneus | 21 | -70 | 19 |  |
| Right Lingual Gyrus | 21 | -70 | -14 | 46 |
| Left Middle Frontal Gyrus | -24 | 8 | 49 | 41 |
| Left Postcentral Gyrus | -27 | -31 | 58 | 26 |
| Left Precentral Gyrus | -30 | -25 | 67 |  |
| Left Supplementary Motor Area | -6 | -13 | 64 | 23 |
| Left Precentral Gyrus | -18 | -13 | 70 |  |
| Right Superior Frontal Gyrus | 24 | -4 | 64 | 14 |
| Right Inferior Frontal Gyrus (p. Opercularis) | 33 | 14 | 34 | 13 |
| Left Medial Temporal Pole | -39 | 14 | -32 | 7 |

MNI coordinates represent the location of the peak voxels. The peak voxels of each cluster with the cluster size are followed by separate maxima (8mm apart) within the cluster.

#### Group by Task interaction effect

We further found a significant group by task interaction, delineating an increased representation of DPA-related WM in older adults’ right inferior temporal gyrus (BA19) and right perirhinal cortex (PRC; BA36), while young adults and synaesthetes showed increased representation of DMS-related WM in PRC (Fig. 3).

**Figure 3.**
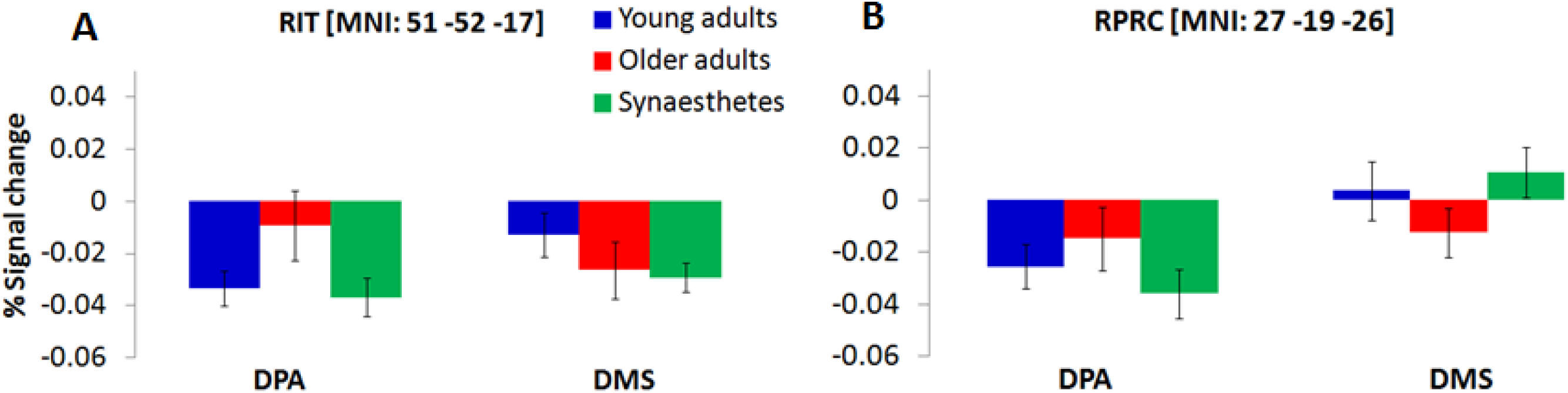
Significant group by task interaction illustrating the WM-type represented in the Right Inferior Temporal Gyrus, RIT (**A**) and Right Perirhinal Cortex, RPRC (**B**) of young adults, older adults and synaesthetes. Percent signal change was extracted from each mean cluster activity and is plotted relative to delay onset. Synaesthetes and young adults exhibited a reduction of BOLD-signal for the maintenance of stimuli that were recalled from memory (DPA-task) in RIT (**A**) and RPRC (**B**). During the DMS-task, the signal reduction was attenuated in synaesthetes and young adults in the RPRC (**B**), but was still present in the RIT (**A**). Error bars represent the standard error of the mean. RIT = Right Inferior Temporal Gyrus, RPRC = Right Perirhinal Cortex.

Separate one-way ANOVAs revealed further group specific effects for each WM task: For the DMS-task (Fig. 4A) we found a significant group effect in the left middle frontal gyrus (BA 9; peak in MNI: -21 8 52), while the more cognitively demanding DPA-task (Fig. 4B) yielded a significant group effect in the left anterior middle frontal gyrus (BA 10; peak in MNI: -30 62 4) and right inferior frontal sulcus (BA44; peak in MNI: 30 14 37). No other group differences were detected. To examine the group differences more closely, we calculated contrast estimates from the mean cluster values of each task and computed Tukey post hoc tests. Older adults showed greater mean activity in frontal regions relative to young adults and synaesthetes in both tasks, as expected. However, while the enhanced activity in the left middle frontal gyrus was non-significantly different in older relative to young adults in the DPA-task (BA10: old > young: p = 0.683) or the DMS-task (BA9: old > young, p = 0.062), the enhanced activation in older adults relative to synaesthetes was always significant (DPA, BA10: p = 0.001; DMS, BA9: p = 0.001). Moreover, for both tasks we found significantly enhanced activity in the left middle frontal gyrus in young adults relative to synaesthetes (DPA, BA10: p = 0.008; DMS, BA9: p = 0.026). These results extend our predictions of enhanced frontal activation in older adults, showing enhanced frontal activation of young and older adults relative to synaesthetes. The only region showing enhanced activity in older adults relative to synaesthetes (p = 0.001) and young adults (p = 0.003) was the right inferior frontal gyrus, while the difference between synaesthetes and young adults in this region was not significant (p = 0.931).

**Figure 4.**
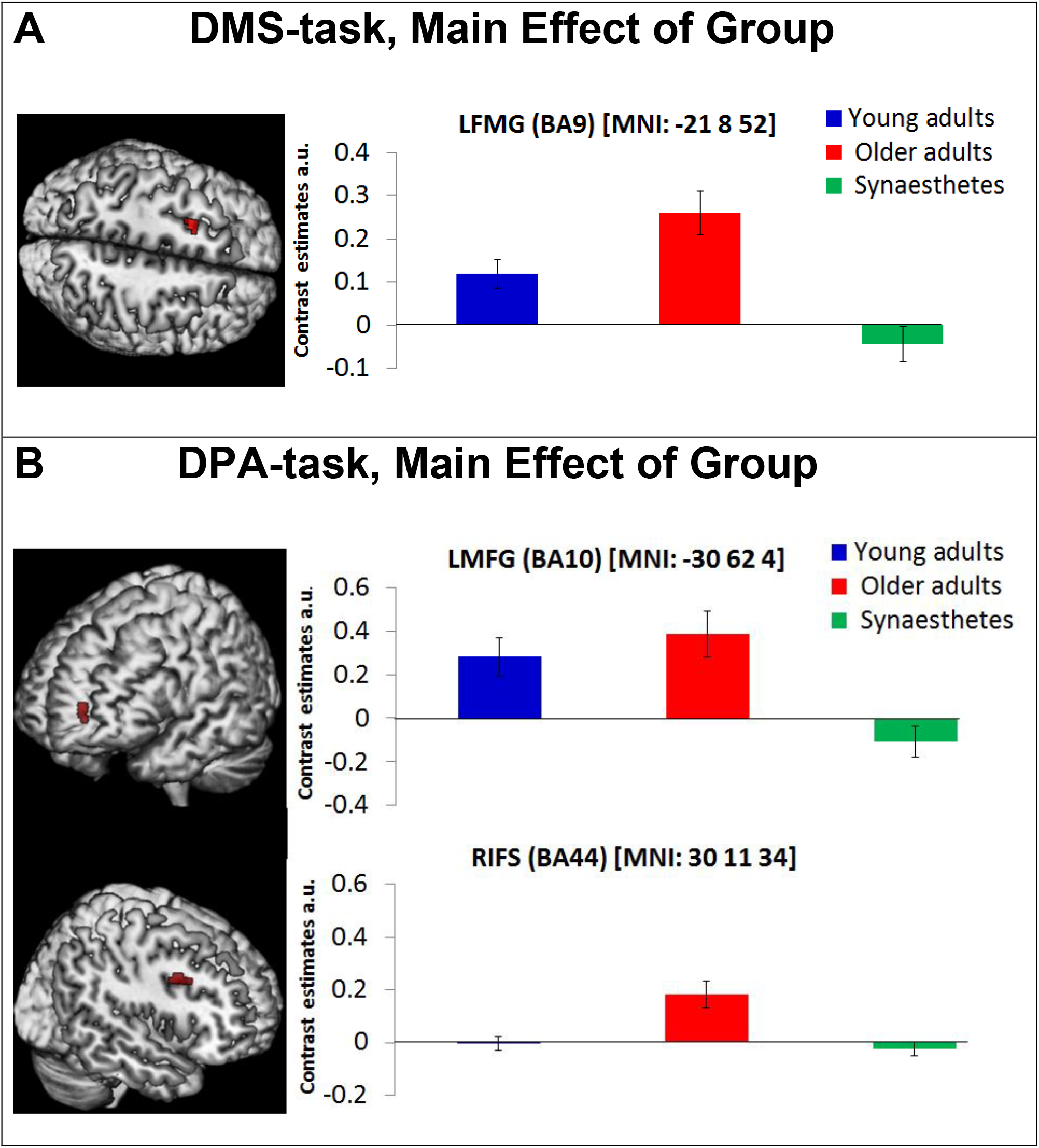
The DMS-task yielded a main effect of group in the Left Middle Frontal Gyrus (BA9) (**A, left**). The DPA-task yielded a main effect of group in the Left Middle Frontal Gyrus (BA10) and the Right Inferior Frontal Gyrus (BA44; **B, left)**. Contrast estimates are reported for each mean cluster activity of DMS and DPA (**A and B, right**). Error bars represent the standard error of the mean. Synaesthetes showed significantly lower activity relative to older adults in all frontal regions. Synaesthetes also exhibited significantly lower activity than young adults in the Left Middle Frontal regions of the DMS and DPA task (**A and B, top**). Young adults showed significantly lower activity than older adults in the RIFS of the DPA-task (**B, bottom**), but not in the Left Middle Frontal regions of the DMS and DPA task (**A and B, top**). LMFG = Left Middle Frontal Gyrus, RIFS = Right Inferior Frontal Sulcus, a.u. = arbitrary unit

## Discussion

The present fMRI study examined whether the disparate sensory-perceptual abilities in old age and grapheme-colour synaesthesia differentially affected brain activity during visual working memory (VWM). Our findings revealed reduced activation in synaesthetes relative to young and older adults across two working memory tasks. These results are in accord with our previous study (Pfeifer et al., 2016) investigating visual associative memory in the same participants and provide evidence for a differentiated visual system supporting higher level cognitive processes (VWM and long term memory) in synaesthesia. Importantly, we show here that this applies to the delay period in working memory (in which the screen was blank) and, hence, group differences cannot be due to visual perception per se but must reflect differences in the maintenance of internal visual representations.

Whole-brain analyses of the present study yielded a group effect in a number of brain regions, notably, the superior and inferior occipital and the inferior temporal gyrus, the precentral and middle frontal gyrus and the medial temporal pole. Extracting the time course of these regions revealed that the synaesthetes’ BOLD signal during the delay period was consistently below young and older adults. Reduced fMRI BOLD signal during delay periods of VWM tasks has been associated with a differentiated neural system that selectively codes for dedicated features (Druzgal and D’Esposito, 2001; Ranganath et al., 2004). During VWM, stimuli are mentally represented in visual cortex and expected after a delay period. The representation and expectation of stimuli ‘sharpens’ receptive units within a well differentiated visual system, resulting in sparse but efficient activation of dedicated neural populations that are manifested as low fMRI BOLD responses (Kok et al., 2012). Having observed reduced BOLD signal in synaesthetes relative to young and older controls in the present WM tasks with achromatic fractal images, our findings suggest that the neural populations supporting these stimuli are more distinctive in synaesthetes compared to the other two groups. Post-hoc tests between young synaesthetes and age-matched young adults further revealed a synaesthesia-specific reduction in BOLD signal, which was exclusively found in posterior visual regions including the inferior occipital and inferior temporal cortex. Our argument is strengthened by our previous study (Pfeifer et al., 2016), in which we analysed the cue and target periods of the same dataset, focusing on cued retrieval and recognition. As in the present study, synaesthetes showed reduced BOLD signal in early visual and inferior temporal cortex relative to young and older adults. However, this finding was limited to the cued retrieval stage and not seen during recognition. Similar to WM, cued retrieval requires an internally directed process to mentally hold, or search for, appropriate stimuli respectively. Thus, the consistently reduced BOLD signal in synaesthetes during VWM and cued retrieval suggests greater efficiency in synaesthetes’ visual cortex and highlights age and individual differences during internally directed cognitive processes. The results of the present study are in line with the sensory recruitment model of VWM (Serences et al., 2009; Lee and Baker 2016), which holds that perceptual stimuli are mentally represented in dedicated, feature selective visual areas. From the model it follows that enhanced perceptual qualities, as in synaesthesia, translate into enhanced VWM, which was evidenced in the present study by the reduced activation in synaesthetes’ visual cortex. Arguably, the reduced visual cortex activation during VWM and cued retrieval could be the result of using the same sample across the current and previous study (Pfeifer et al., 2016). However, the fact that synaesthetes demonstrated greater visual cortex activity relative to young and older adults during recognition (Pfeifer et al., 2016) shows that our sample of synaesthetes did not chronically express lower activation across all cognitive processes. Rather, synaesthesia and age-related perceptual mechanisms contributed differently to bottom-up and top-down processes (recognition, and WM and retrieval, respectively).

A repeated finding in our time course analysis was a negative BOLD response during the delay period in synaesthetes’ bilateral inferior and superior occipital cortex, and, most prominently, in the left medial temporal pole (LMTP) (Figure 2). The negative BOLD response has been interpreted as a diversion of blood flow to active brain areas, causing a local dip in non-used regions (Wade, 2002). For example, Amedi et al. (2005) found that the amplitude of the negative BOLD response in auditory cortex correlated with a positive BOLD signal in visual cortex during a visual imagery task. The results suggested that blood flow was diverted from an unused sensory region (i.e. auditory cortex) to relevant areas in visual cortex to engage in visual imagery. A similar interpretation might explain the negative BOLD response within our synaesthetes’ LMTP. The temporal pole is involved in semantic processing (Lambon Ralph et al., 2009) and object naming (Price et al., 2005). In the present study, naming abstract fractals might have assisted with the sustained representation of the images in WM. However, given that synaesthetes perform well on visual imagery (Barnett and Newell, 2007; Spiller et al., 2015), it is plausible that they used a purely visual strategy, thus causing the negative BOLD response in the LMTP that constituted an irrelevant brain region. The interpretation of diverted blood flow falls short, however, in explaining the negative BOLD response observed in synaesthetes’ posterior occipital regions that were actively engaged during our VWM task. Bressler et al. (2007) offer an alternative explanation, suggesting that the negative BOLD signal, particularly in visual cortex, carries relevant and stimulus-specific information. Their interpretation converges with fMRI studies using multivoxel pattern analysis (MVPA), showing that negative BOLD signal in visual regions carries content-specific information of perceptually encoded stimuli during mental representation (Serences et al., 2009; Riggall and Postle, 2012). Hence, it is possible that the role of the negative BOLD signal observed in synaesthetes is to carry information content in posterior visual regions, but not in anterior semantic processing areas such as the LMTP.

Concerning aging, our results are consistent with previous reports and suggest age-related dedifferentiation in posterior visual regions, concomitant with increased compensatory top-down functions from PFC. Relative to young adults and synaesthetes, older adults showed the highest mean amplitude in all occipito-temporal areas during VWM (Figure 2). Previous research has shown that neural populations in occipito-temporal regions become less distinctive with age and are characterised by non-specific activation of feature-selective areas during perception (Park et al., 2004; 2012) and visual imagery (Kalkstein et al., 2011). For example, Kalkstein et al. (2011) found that visual imagery of cued faces and moving dots activated respective face and motion selective regions in young adults, while older adults recruited both areas irrespective of the imagined stimulus type. Although the mental representation of fractal stimuli in our study did not target feature-selective regions as in the study by Kalkstein et al. (2011), the enhanced activation of older adults’ early visual regions relative to young adults and synaesthetes suggests activation of broad, non-specific neural ensembles, which manifested as increased fMRI BOLD signal in superior and inferior occipito-temporal regions.

Further evidence for age-related dedifferentiation was observed by a significant group by task interaction in the right inferior temporal cortex (RIT) and the right perirhinal cortex (RPRC). Consistent with the notion that older adults show greater activity as a result of neural broadening, we found less signal reduction (i.e. higher mean amplitude) in RIT and RPRC for older adults relative to young adults and synaesthetes during DPA-related WM (Figure 3). By contrast, young adults and synaesthetes exhibited greater activity relative to older adults in RPRC during DMS-related WM. The PRC is sensitive to minimal feature changes in complex visual stimuli (Gonsalves et al., 2005; Henson et al., 2003). Our DMS task required precise feature mappings of stimulus representations held in WM with a subsequently appearing perceptual target. Young adults and synaesthetes might have benefitted from recruiting the PRC to resolve any feature ambiguity in the DMS task, while older adults relied significantly less on this region, possibly as a result of age-related decline in PRC function (Ryan et al., 2012).

Consistent with our prediction, we observed enhanced PFC activity in older adults relative to young adults and synaesthetes across both VWM tasks (Table 4). Enhanced activity in PFC serves as a compensatory strategy for age-related neural dedifferentiation in posterior visual regions and has been described as the posterior-to-anterior shift in aging (PASA; Davis et al., 2008). Evidence for the PASA account comes from studies showing that age-related increases in PFC correlated with behavioural accuracy (Davis et al., 2008) and reduced activity in occipito-temporal regions (Cabeza et al., 2004; Davis et al., 2008). The behavioural accuracy scores in our paradigm were high across the two WM tasks and comparable between groups (no main effect of group was found) (Table 2). Thus, the enhanced frontal activation in older adults can be interpreted as a compensatory strategy to achieve comparable performance to young adults and synaesthetes. The fact that we only included successful and confident trials in our fMRI analysis adds further confidence to the older adults’ compensatory mechanisms via PFC to achieve accurate and confident performance scores.

Of further interest to the present study were the modulatory influences of task difficulty on brain activation. The results of our two WM-tasks, DMS (maintenance of cued images, low WM-load) and DPA (maintenance of retrieved images from memory, high WM-load) demonstrated that the neural correlates of WM are task-dependent: The contrast DPA > DMS activated the left lateral PFC and superior parietal cortex, as would be expected from a retrieval-related WM task (Vilberg and Rugg, 2012). By contrast, DMS > DPA activated the medial PFC, lateral and medial temporal regions and inferior parietal cortex, which is consistent with a pure visual working memory task (Ranganath, 2006).

Separate analyses for the DMS and DPA task further elucidated age and individual differences in compensatory strategies employed via the PFC. We observed reduced activity in PFC in synaesthetes relative to young and older adults in both WM tasks, suggesting that synaesthetes required less compensatory mechanisms than the other two groups. Moreover, the group differences found in PFC reflected the specific type of WM: the DMS-task yielded a significant group effect in the left middle frontal gyrus (BA 9), which is classically associated with the maintenance of information in WM, including the reactivation of just-seen, transiently stored material (Curtis and D’Esposito, 2003; Raye et al., 2002). Older adults, who activated this region more strongly than synaesthetes (p=0.001) and young adults (p=0.062), might have compensated for behavioural performance, which did not differ between groups. The fact that young adults also showed significantly enhanced activity relative to synaesthetes (p = 0.026) highlights the effect of synaesthesia, indicating greater WM-related efficiency in synaesthetes that is less dependent on top-down control mechanisms. Our findings are in line with the distributed model of WM that considers collaborative operations of PFC and posterior visual regions (Postle, 2006; D’Esposito and Postle 2015; Lee and Baker 2016), suggesting that top-down control from PFC is alleviated in cases where the neural sensitivity in posterior visual regions is enhanced (as in synaesthesia).

The DPA-task yielded two group effects, one in the right inferior frontal sulcus and another in the left middle frontal gyrus, corresponding to the lateral region of BA10. We attribute the group differences in the right inferior frontal sulcus to a specific age-related dedifferentiation, given the enhanced activity in older adults relative to both, young adults and synaesthetes. Specifically, aging has been associated with a hemispheric asymmetry, whereby older adults show less left-lateralized activity than young adults and often activate additional right frontal regions (Cabeza, 2002), consistent with our finding. Our behavioural results shed further light on the observed age-related compensatory mechanisms. Older adults performed significantly poorer than the other two groups on the dissimilar condition of the DPA task. Since our fMRI analyses were carried out using trials from the dissimilar condition only (constituting high retrieval and WM-load), our imaging results converge with behavioural findings and demonstrate that age-related compensatory mechanisms via PFC increase with task difficulty.

The group effect in BA10, which has been associated with the recollection of contextual details in associative memory tests (Simons et al., 2005a; Simons et al., 2005b), reflects memory-related processing differences inherent in the DPA-task. Although the instruction was to use the cue for retrieval and the delay for maintaining the retrieved pair-associates, it is likely that some participants continued to re-activate the to-be-maintained information during the delay-period. In this sense, the group differences found in lateral BA10 reveal the additional memory demands imposed by DPA-related over DMS-related WM. Interestingly, young and older adults showed significantly enhanced activity in BA10 relative to synaesthetes, suggesting that it was the specific retrieval-related maintenance subserved by this region during which synaesthetes demonstrated greater efficiency.

In conclusion, our data suggest a differentiated visual system in synaesthetes relative to young and older adults that supports VWM maintenance of non-synaesthesia inducing stimuli. The enhanced cortical sensitivity in synaesthetes’ visual areas (Barnett et al., 2008; Terhune et al., 2011) was reflected by reduced occipito-temporal activation during VWM and is consistent with the sensory recruitment model (Serences et al., 2009; Lee and Baker 2016). Beyond sensory recruitment, synaesthetes also showed diminished top-down activation from PFC across two WM tasks that varied in cognitive demand. This finding dovetails with the suggested frontal control functions over posterior regions as envisaged by distributed models of WM (Postle, 2006; D’Esposito and Postle 2015; Lee and Baker 2016) and lends support for an overall neural efficiency of synaesthetes’ brains to assist VWM maintenance. The novelty of our finding is that sensory-perceptual processing differences inherent in the three groups translated into differences in VWM processing. Importantly, this finding converges with, and extends our previous result for visual associative memory (Pfeifer et al. 2016), demonstrating the utility of multiverse analyses of the same dataset to obtain a holistic picture across different cognitive processes (Steegen et al., 2016). Specifically, the synaesthetes’ reduced BOLD signal in visual cortex during VWM (present study) and cued retrieval (Pfeifer et al., 2016) suggests that internally directed cognitive processes recruit selective neural populations, which manifest as reduced fMRI activation in a sensitive visual system (as in synaesthesia). To our knowledge this is the first evidence from aging and synaesthesia that links bottom-up perceptual qualities with top-down WM maintenance, and is in support of the sensory recruitment and distributed models of working memory.

